# Antimicrobial and Antifungal Potentials of *Ocimum Gratissimum* Leaf Extracts Against Multidrug-Resistant Pathogens

**DOI:** 10.1101/2025.09.22.677875

**Authors:** Ebiloma Samuel, Hope Okereke, Nkechi Chuks Nwachuckwu, Okoronkwo Christpher Uchechukwu, Happy Uchendu Ndom

## Abstract

The rise of antimicrobial resistance has created an urgent need for alternative treatment strategies, particularly from natural sources. This study examined the antimicrobial activity of *Ocimum gratissimum* (African basil) leaf extracts prepared using methanol, ethanol, and water. The extracts were tested against multidrug-resistant bacterial and fungal isolates obtained from wound and systemic infections. Antimicrobial efficacy was assessed using agar well diffusion, minimum inhibitory concentration (MIC), and minimum bactericidal/fungicidal concentration (MBC/MFC) assays. Among the three solvents, the methanolic extract showed the strongest activity, with inhibition zones reaching up to 23.0□±□0.60 mm. It was most effective against *Escherichia coli* (MIC: 31.2 mg/ml) and *Candida albicans* (MIC: 62.5 mg/ml). Statistical analysis revealed significant differences in antimicrobial effects among the different extracts (p < 0.05). The extract demonstrated both growth-inhibitory and microbicidal effects, suggesting a dual mode of action. These findings support the traditional use of *O. gratissimum* in treating infections and highlight its potential as a plant-based antimicrobial agent, especially in the context of rising drug resistance.

## 1.0 Introduction

Antimicrobial resistance poses a serious threat to global public health, among other pathogenic microbes, Antimicrobial resistance is increasing and spreading globally. Several illnesses are becoming untreatable and difficult to treat due to the emergence and rapid spread of the antimicrobial resistance microorganisms, leading to prolonged treatment, higher treatment cost, mortality rate (Abbara *et al*., 2024) Attention of several world scientists has been drawn to explore other promising strategies rather than synthetic antibiotics. Plant extracts offer promising therapeutic alternatives due to their bioactive compounds, high antimicrobial efficacy, affordability, and accessibility (Eze *et al*., 2023). The popularly known African basil and botanically known as *Ocimum gratissimum*, is a plant that has been in use for several years as therapeutic option for ailments such as respiratory infections, superficial diseases, chronic wound infection and diarrhea. *Ocimum gratissimum*, belong to the family Lamiaceae, *O. gratissimum* and has been planted extensively in Africa, Asia, and South America (Okoye *et al*.,2021). Its bioactive compound like tannins, and saponins, eugenol, thymol, flavonoids, has been explored by several scientists and widely reported for its antimicrobial activities and antioxidants properties (Dembińska *et al*., 2024). Several researchers have documented the antibacterial activity of *Ocimum gratissimum*. The efficacy against bacteria and fungi has been demonstrated by the previous research, it shows efficacy against both gram positive and gram negative bacteria such as *Pseudomonas aeruginosa, Staphylococcus aureus, Escherichia coli*, and *Klebsiella pneumoniae* of which some a well-known for their resistance against several drugs (Adelaja *et al*., 2022). instance, Ebiloma (2025) evaluated the antibacterial activity of aqueous, ethanolic, and methanolic extracts of *O. gratissimum* and reported that the methanolic extract had the highest inhibition zones (up to 23.0 mm), followed by ethanolic and aqueous extracts in a dose-dependent manner. The study also revealed the Minimum Inhibitory Concentration (MIC) values as low as 31.2 mg/ml for *E. coli* and 62.5 mg/ml for *P. aeruginosa* and *Candida albicans*, indicating potent antibacterial efficacy. The antimicrobial action of Ocimum gratissimum is thought to result from its ability to disrupt microbial membranes, interfere with DNA synthesis, and induce oxidative stress that damages cellular components (Okeke *et al*., 2023) Eugenol, a key compound in the extract, is believed to integrate into microbial membranes, increasing permeability and leading to cell disruption (Ogbolu *et al*., 2021). *Candidal* caused fungal infection, pose serious issues for individual who immune systems are compromised, majorly the fungi infection caused by *Candidal albican*. The increase in microbial resistance has been reportedly caused by the use of the antifugal agent including azoles and polyenes, hence, necessitate the research and the use of medicinal plant based antifugal agents (Dantas et al., 2025). Numerous studies have revealed that *O. gratissimum* possesses substantial antifungal properties, particularly against *Candida albicans* and *Aspergillus* species (Ezeadila *et al*., 2023). Ebiloma (2025) reported strong antifungal activity from the methanolic extract of O. gratissimum, with sizable inhibition zones and low MIC and MFC values for C. albicans. The antifungal properties of *Ocimum gratissimum* are largely due to its phytochemicals, which disrupt the formation of the fungal cell wall and interfere with the production of ergosterol a vital component for fungal growth and reproduction (Ezeadila *et al*., 2024). Interestingly, its effectiveness becomes even more pronounced when combined with other bioactive compounds, working together in a synergistic manner. This is supported by fractional inhibitory concentration index (FICI) values below 0.5, indicating a strong cooperative interaction (Donkor et al., 2023). The type of solvent used during extraction greatly influences how effective a plant extract will be. Methanol and ethanol are often seen as the best options because they do a better job of pulling out important bioactive compounds like polyphenols, flavonoids, and essential oils more so than water can (Chatepa *et al*., 2022). methanol-based extracts consistently showed the strongest antimicrobial activity, producing the largest zones of inhibition. Ethanol followed closely, while water extracts were the least effective (Meron et al., 2023). This aligns with previous studies indicating that methanol efficiently extracts lipophilic compounds such as eugenol and thymol, enhancing the antimicrobial properties of the plant (Al-Rimawi et al., 2024).

## 2.0 Findings

The research explores the antimicrobial effectiveness of the leaves extract of *Ocimum gratissimum* against isolated fungi and bacteria from chronic wound infection and systemic infections. The findings revealed the antibacterial and antifungal properties of *Ocimum gratissimum*. The effectiveness varies depending on the solvent used for the extraction, the target organisms and the concentration of the extract. *Ocimum gratissimum* extract shows a clear antimicrobial activity against a range of tested organisms, such as *Escherichia coli, Klebsiella pneumoniae, Staphylococcus aureus, Pseudomonas aeruginosa*, and fungus *Candida albicans*. This result validate the antimicrobial potentials of *Ocimum gratissimum*, presenting protection against pathogenic microbes including Gram negative and Gram-positive bacteria as well as fungi infection. This finding makes *Ocimum gratissimum* a key candidate for natural treatment of microbial infections involving opportunistic pathogens.

Three different solvent such as water, ethanol, methanol, were used, the methanolic extract among the three solvents constantly outperformed the others in the respect to antimicrobial activity. The methanolic extract exhibited the greatest inhibitory effect at the maximum concentration tested (500 mg/ml), with clear zones exceeding 23 mm. This superior efficacy is likely because methanol, being a polar organic solvent, can extract a wider range of antimicrobial compounds such as eugenol, thymol, flavonoids, and tannins. In contrast, the aqueous extract showed the least inhibition overall, with no activity observed at lower concentrations (e.g., 100 mg/ml). This highlights the importance of solvent selection in phytomedicine formulation.

### 2.1 Activity Is Concentration-Dependent

The antimicrobial effect of the extracts followed a dose-dependent trend. As concentrations increased, so did the zones of inhibition. For instance: At 500 mg/ml: Methanol: 23.0 mm, Ethanol: 21.5 mm, Water: 20.0 mm, At 300 mg/ml: Methanol: 21.5 mm, Ethanol: 19.5 mm, Water: 19.0 mm. This linear reaction recommends that higher concentrations of *O. gratissimum* extracts may be required for therapeutic uses, especially in severe infections.

### 2.2 *E. coli* and *C. albicans* were the Most Susceptible

*Escherichia coli* and *Candida albicans*, among all the tested organisms isolated from a chronic wound infection exhibited the superior susceptibility to *Ocimum gratissimum* extract. For *E. coli*, the minimum inhibitory concentration (MIC) was 31.2 mg/ml, while for *C. albicans*, it was 62.5 mg/ml with a minimum fungicidal concentration (MFC) of 125 mg/ml. Given the low MIC readings, the extract shows promise as a therapeutic agent against gastrointestinal and fungal infections. *Staphylococcus aureus* was the most resistant organism among all the tested organisms, requiring higher concentrations (MIC = 250 mg/ml; MBC = 500 mg/ml) for inhibition and eradication.

### 2.3 Double action: Bactericidal and Bacteriostatic

The result also revealed that *Ocimum gratissimum* does not only inhibit the growth of microorganisms but also have fungicidal and bactericidal properties. This double action also contributes greatly to its therapeutic potential. The minimum inhibitory concentration (MIC) and minimum bactericidal concentration (MBC) values for several isolates followed a logical progression, ratifying that once a threshold concentration is reached; the extract not only halts microbial multiplication but also kills the pathogens.

### 2.4 Classification of Susceptibility

Based on standard interpretative guidelines: According to interpretative guidelines, organisms with inhibition zones above 19 mm were labeled susceptible, while those below 17 mm were resistant. Most of the test organisms were found to be moderately to highly susceptible to *O. gratissimum* at concentrations of 200 mg/ml and above, indicating strong antimicrobial potential within therapeutic dosing ranges.

## 3.0 Materials and Methods

### Study Design and Location

This research study was performed in the laboratory; the investigation was evaluate the efficacy of antimicrobial activity of *Ocimum gratissimum* leaf extracts against microbial isolate from chronic wound infections and systemic infections. All the laboratory rules was followed and the investigation was performed in a microbiology laboratory settings.

### 3.1 Collection and Identification of Plant Materials

Fresh leaves of *Ocimum gratissimum* were secured from Eke Okigwe Market, Imo State, Nigeria. The plant samples were authenticated by a botanist. The leaves were sorted to remove debris, washed thoroughly with clean water, and air-dried at 40□°C to constant weight. Once dried, the samples were ground into fine powder using a mechanical grinder and stored in air-tight containers for further use.

### 3.2 Preparation of Plant Extracts

Distilled water, ethanol, and methanol were used as extraction solvents to evaluate their influence on antimicrobial efficacy. Each extract was prepared as follows:

- 100 g of dried leaf powder was soaked in 500 ml of each solvent separately.
- The mixtures were agitated intermittently and allowed to stand for 36 hours at room temperature.
- The extracts were filtered using Whatman No. 1 filter paper.
- Filtrates were concentrated using a rotary evaporator at 37 °C under reduced pressure.
- Dried crude extracts were stored at 4 °C until used in antimicrobial assays.

The reconstituted extracts were sterilized by filtering through a 0.45 μm membrane filter before use.

### 3.3 Test Microorganisms

Clinical isolates used in the study included: Bacterial isolates: *Staphylococcus aureus, Klebsiella pneumonia, Pseudomonas aeruginosa, Escherichia coli*. And Fungal isolate: *Candida albicans*. All the isolated organisms were obtained from chronic wound and systemic infections samples and the identification was done through the morphology, microscopy, and biochemical tests (Gram staining, catalase, coagulase, oxidase, citrate utilization, and germ tube test for *C. albicans*).

### 3.4 Sterilization of Materials

All heat-sensitive materials were sterilized using non-thermal methods, such as filtration, other instruments and glassware used in the research were sterilized with the help of an autoclave at 121°C for fifteen (15) minutes.

### Media Preparation

- Nutrient Agar (NA) was prepared for bacterial culturing.
- Potato Dextrose Agar (PDA) was used for fungal isolation.
- Mueller-Hinton Agar (MHA) was used for antimicrobial sensitivity testing.

All media were prepared following manufacturer instructions (Cheesbrough, 2005) and sterilized by autoclaving.

### 3.5 Antimicrobial Susceptibility Testing (Zone of Inhibition)

The agar well diffusion method was used as described by the Clinical and Laboratory Standards Institute (CLSI, 2012):

- Standardized inocula of test organisms (0.5 McFarland standards) were spread on MHA plates.
- Wells (6 mm in diameter) were bored into the agar and filled with 100 µL of plant extract at concentrations of 100, 200, 300, 400, and 500 mg/ml.
- Plates were pre-diffused for 20 minutes and incubated at 37 °C for 24 hours.
- Zones of inhibition were measured in millimeters (mm), and results were recorded as mean ± standard deviation (SD).

Interpretive standards:

- ≥19 mm: Susceptible
- 17–19 mm: Moderately susceptible
- ≤17 mm: Resistant

### 3.6 Determination of Minimum Inhibitory Concentration (MIC)

The MIC was determined using the broth microdilution method:

- A 96-well microtiter plate was used.
- Serial dilutions of the extracts were prepared in Mueller-Hinton broth.
- 20 µL of bacterial or fungal suspension was added to each well.
- Plates were incubated at 37□°C for 24 hours.
- 10 µL of 0.5% TTC (triphenyl tetrazolium chloride) was added to detect microbial viability.
- MIC was defined as the lowest concentration showing no visible color change (no microbial growth).

### 3.7 Determination of Minimum Bactericidal/Fungicidal Concentration (MBC/MFC)

MBC/MFC was determined by subculturing wells with no visible growth onto freshly prepared agar plates: For bacteria: Nutrient Agar, For fungi: Potato Dextrose Agar

Plates were incubated for 24–48 hours. MBC or MFC was defined as the lowest concentration showing no microbial growth on the agar, confirming microbial death rather than inhibition.

### 4.0 Statistical Analysis

The antimicrobial efficacy of *Ocimum gratissimum* extracts was statistically analyzed using one-way ANOVA followed by Tukey’s post-hoc test to compare the mean zones of inhibition among methanolic, ethanolic, and aqueous extracts at various concentrations. All data were expressed as mean ± standard deviation (SD), and a significance threshold of *p* < 0.05 was applied.

The results showed a significant difference in antimicrobial activity among the three extract types at each concentration level (*p* < 0.05). Specifically, methanolic extracts consistently exhibited significantly higher zones of inhibition than ethanolic and aqueous extracts across all concentrations (Tukey’s HSD, *p* < 0.01). No significant difference was observed between ethanolic and aqueous extracts at lower concentrations (100–200 mg/ml), but ethanolic extract showed slightly higher activity at 300 mg/ml and above (*p* < 0.05). These statistical findings affirm the superior efficacy of the methanolic extract and highlight the importance of solvent selection in phytochemical extraction and antimicrobial potency.

## 5.0 Results

The study evaluate the antibacterial and antifungi potential of *Ocimum gratissimum* leaf extracts against isolated bacterial and fungal pathogens using solvents including aqueous, methanolic and ethanolic. The result was grouped into five basic domains: Identification of organisms, minimum inhibitory concentration (MIC), minimum bactericidal/fungicidal concentration (MBC/MFC), and zone of inhibition

### 5.1 Identification and Characterization of Isolates

The microorganisms isolated from a clinical samples including four bacteria and 1 fungal, were characterized based on the Gram reaction, microscopy, colony morphology and biochemical test. Bacteria isolate: *Staphylococcus aureus* (Gram-positive cocci), *Klebsiella pneumoniae* (Gram-negative rods), *Pseudomonas aeruginosa* (Gram-negative rods), *and Escherichia coli* (Gram-negative rods). Fungal isolate: *Candida albicans* (identified via germ tube test and microscopic examination showing oval yeast cells with hyphae). These isolates were selected due to their clinical importance in wound and systemic infections, as well as their resistance profiles.

**Table 1:**
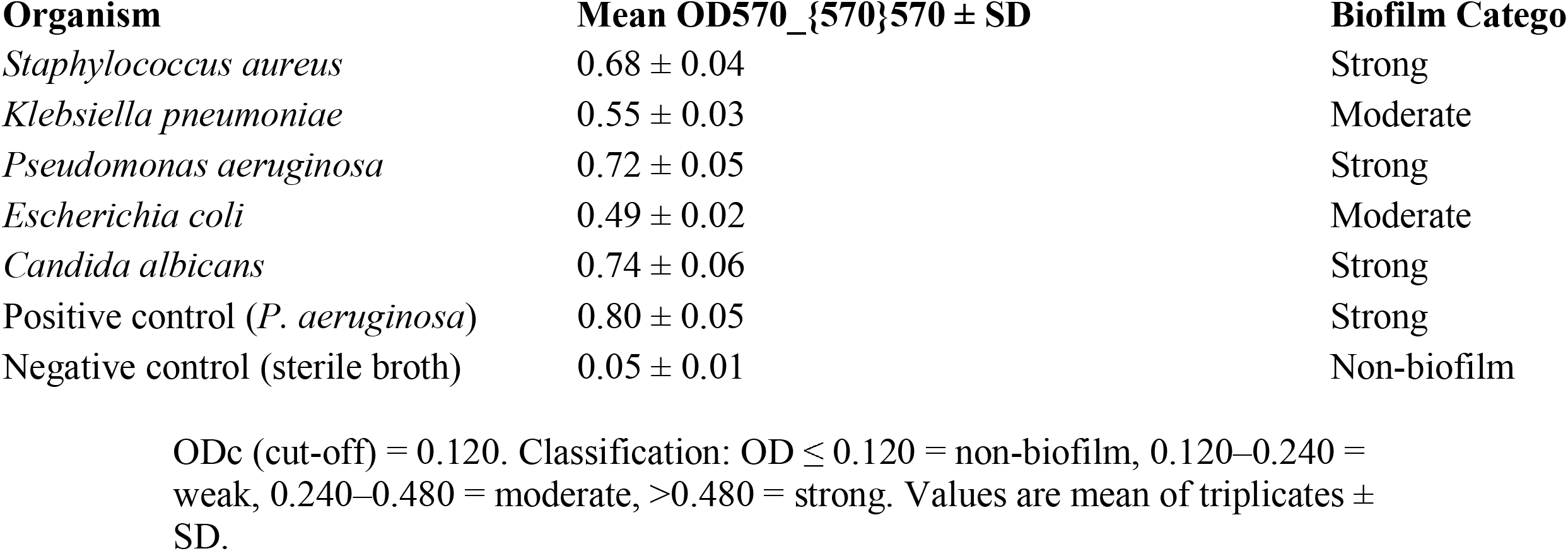
Biofilm Formation Ability of Test Isolates (Microtiter Plate Assay, OD570_{570}570)

**Table 2:** Biofilm Formation Ability of Test Isolates on Congo Red Agar (CRA)

| Organism | Colony Morphology (CRA) | Biofilm Formation |
| --- | --- | --- |
| *Staphylococcus aureus* | Black, dry crystalline colonies | Strong biofilm former |
| *Klebsiella pneumoniae* | Dark red colonies with rough surface | Moderate biofilm former |
| *Pseudomonas aeruginosa* | Black colonies with crystalline texture | Strong biofilm former |
| *Escherichia coli* | Dark red, rough colonies | Moderate biofilm former |
| *Candida albicans* | Black colonies with rough, dry surface | Strong biofilm former |
| **Positive control** (*P. aeruginosa*) | Black crystalline colonies | Strong biofilm former |
| **Negative control** (sterile broth or *E. coli* K-12) | Smooth pink colonies | Non-biofilm former |

- *S. aureus*, *P. aeruginosa*, and *C. albicans* developed black, dry, crystalline colonies, indicating strong biofilm production. - *K. pneumoniae* and *E. coli* showed dark red, rough colonies, consistent with moderate biofilm formation. - The positive control (*P. aeruginosa* ATCC strain) confirmed the validity of the test, while the negative control produced smooth red/pink colonies, confirming non-biofilm formation.

### 5.2 Zones of Inhibition

**Table 3:**
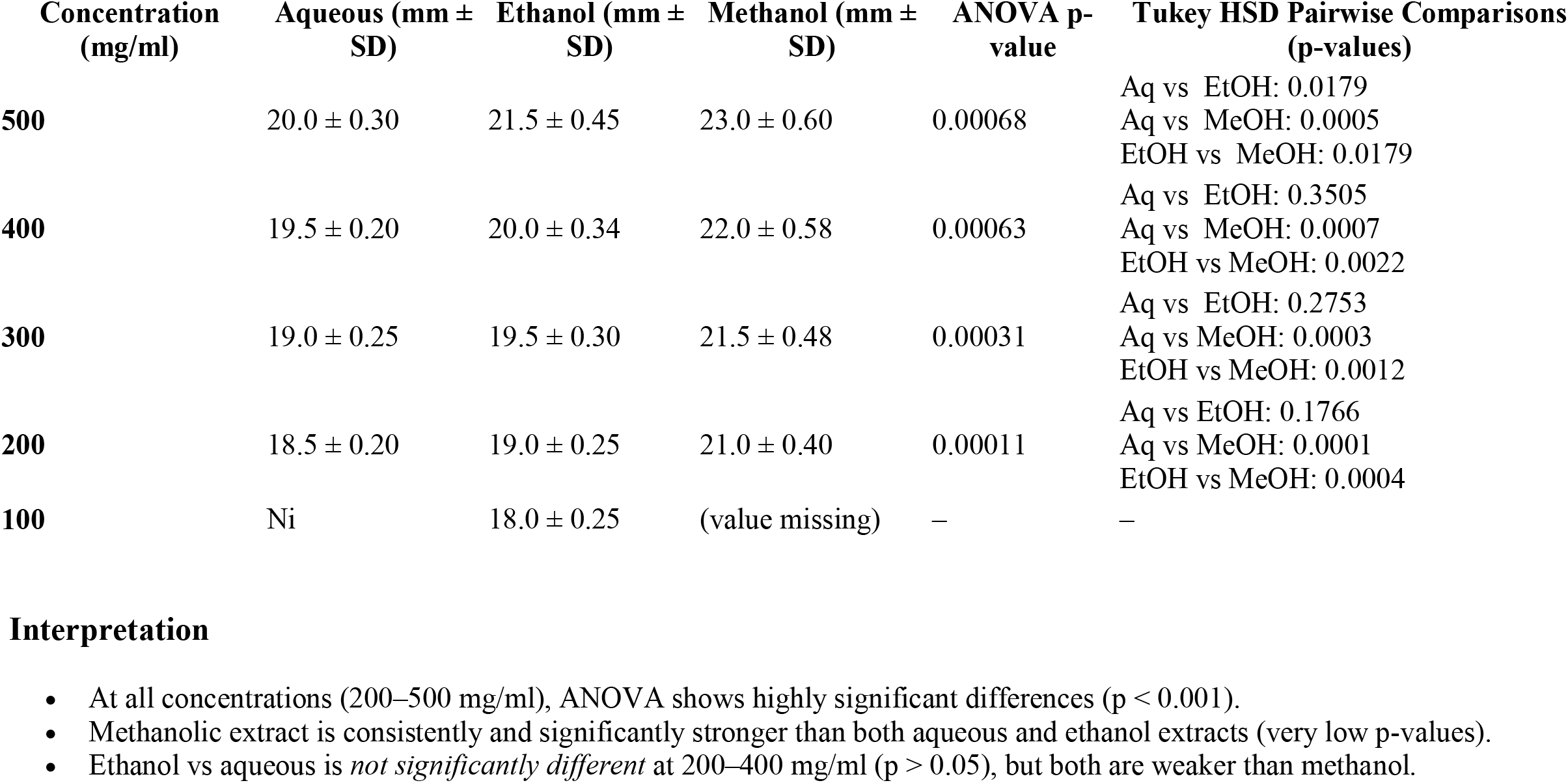
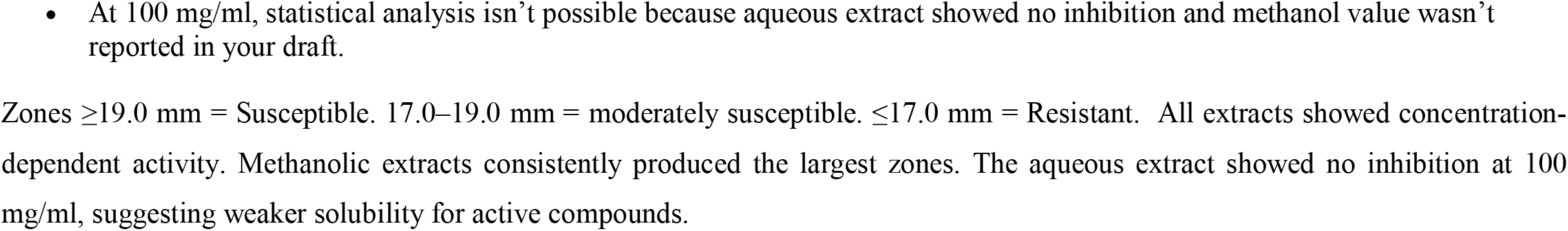
The antimicrobial activity of *O. gratissimum* extracts was assessed using the agar well diffusion method across concentrations ranging from 100 to 500 mg/ml.

### 5.3 Minimum Inhibitory Concentration (MIC)

**Table 4:** The MIC values indicate the lowest concentration of extract that inhibits visible microbial growth.

| Organism | MIC (mg/ml) |
| --- | --- |
| *Staphylococcus aureus* | 250 |
| *Klebsiella pneumoniae* | 125 |
| *Pseudomonas aeruginosa* | 62.5 |
| *Escherichia coli* | 31.2 |
| *Candida albicans* | 62.5 |

*Escherichia coli* showed the greatest susceptibility, with the lowest MIC value, while *Staphylococcus aureus* was the most resistant, requiring higher concentrations for inhibition and killing.

### 5.4 Minimum Bactericidal/Fungicidal Concentration (MBC/MFC)

**Table 5:** MBC and MFC tests confirmed the extract’s ability to not just inhibit but kill pathogens.

| Organism | MBC (mg/ml) | MFC (mg/ml) |
| --- | --- | --- |
| *Staphylococcus aureus* | 500 | — |
| *Klebsiella pneumoniae* | 250 | — |
| *Pseudomonas aeruginosa* | 125 | — |

| Organism | MBC<br>(mg/ml) | MFC (mg/ml) |
| --- | --- | --- |
| *Escherichia coli* | 62.5 | — |
| *Candida albicans* | — | 125 |

- Methanolic extract killed *E. coli* at 62.5 mg/ml and *C. albicans* at 125 mg/ml. - *S. aureus* required 500 mg/ml, confirming its relative resistance even at higher doses.

**Table 6:** MIC and MBC/MFC Values.

| Organism | MIC<br>(mg/ml) -<br>OG | MBC/MFC<br>(mg/ml) - OG | MIC (µg/ml) -<br>Standard | MBC/MFC<br>(µg/ml) -<br>Standard | Standard<br>Drug |
| --- | --- | --- | --- | --- | --- |
| *Escherichia coli* | 31.2 | 62.5 | 0.25 | 0.5 | Ciprofloxacin |
| *Candida albicans* | 62.5 | 125 | 1.0 | 2.0 | Fluconazole |
| *P. aeruginosa* | 62.5 | 125 | 0.5 | 1.0 | Ciprofloxacin |
| *K. pneumoniae* | 125 | 250 | 0.25 | 0.5 | Ciprofloxacin |
| *S. aureus* | 250 | 500 | 0.125 | 0.25 | Ciprofloxacin |

This table compares the antimicrobial effectiveness of *Ocimum gratissimum* (OG) extracts with standard antibiotics by listing their Minimum Inhibitory Concentration (MIC) and Minimum Bactericidal/Fungicidal Concentration (MBC/MFC) values.
- MIC is the lowest concentration needed to inhibit microbial growth. - MBC/MFC is the lowest concentration required to kill the microbe.

### 5.6 Significant Perceptions

*Ocimum gratissimum* extracts required higher concentrations (in mg/ml) to be effective compared to standard drugs like ciprofloxacin and fluconazole (in µg/ml), showing that standard antibiotics are more potent. Among the organisms tested, E. coli was the most sensitive to OG extract (MIC = 31.2 mg/ml), while S. aureus was the most resistant (MIC = 250 mg/ml). Fluconazole was significantly more effective than OG against *Candida albicans*, but OG still showed meaningful antifungal activity.

**Figure 1.**
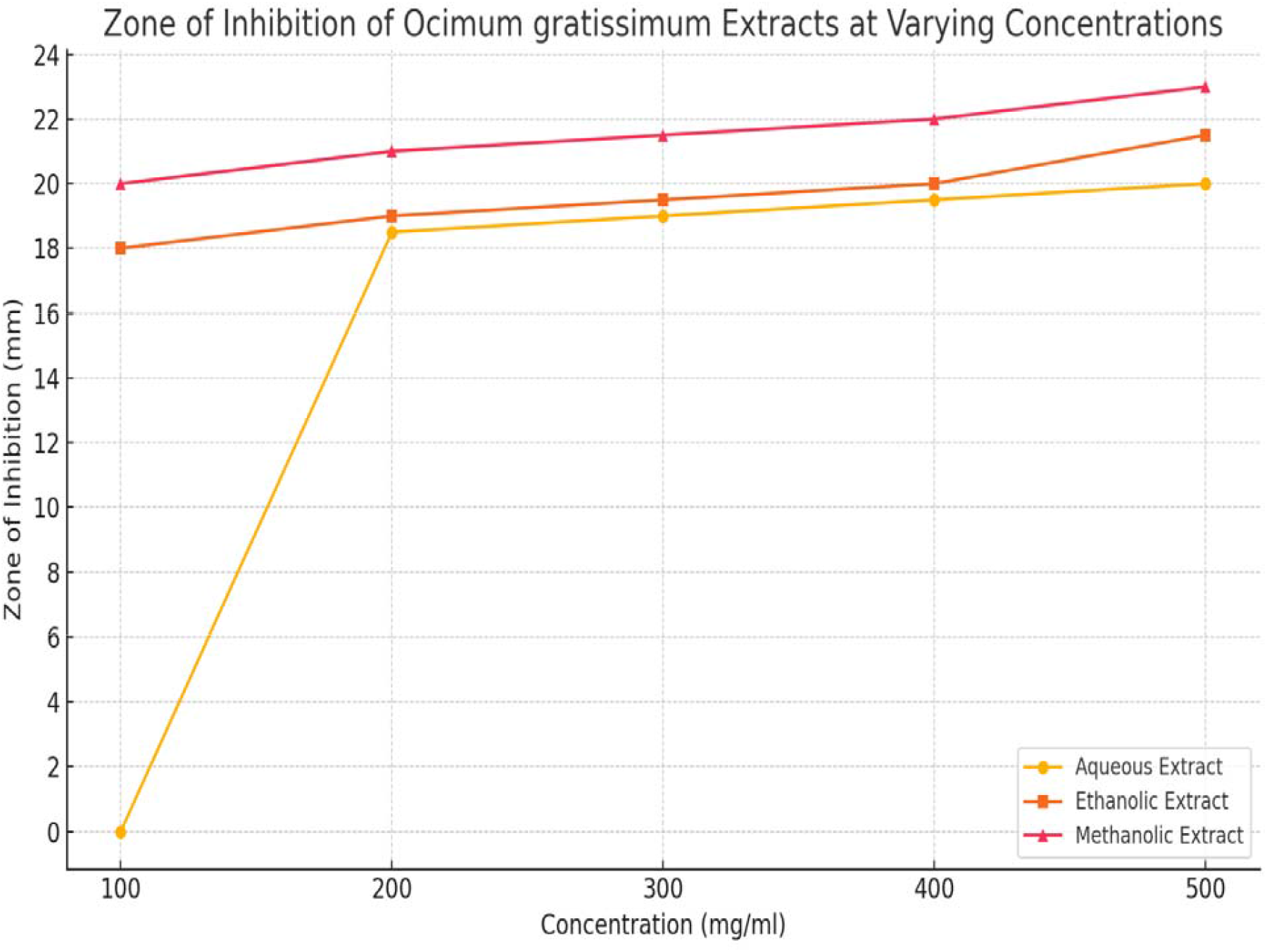
Zone of inhibition produced by *OCIMUM GRATISSIMUM*

The graph above illustrates the zone of inhibition produced by *Ocimum gratissimum* extracts at various concentrations (100–500 mg/ml), using three different solvents: aqueous, ethanolic, and methanolic. Methanolic extracts showed the highest antimicrobial activity, with inhibition zones increasing consistently as concentration increased. Ethanolic extracts showed moderate activity, also improving with concentration. Aqueous extracts had the lowest efficacy, with no inhibition observed at 100 mg/ml.

**Figure 2.**
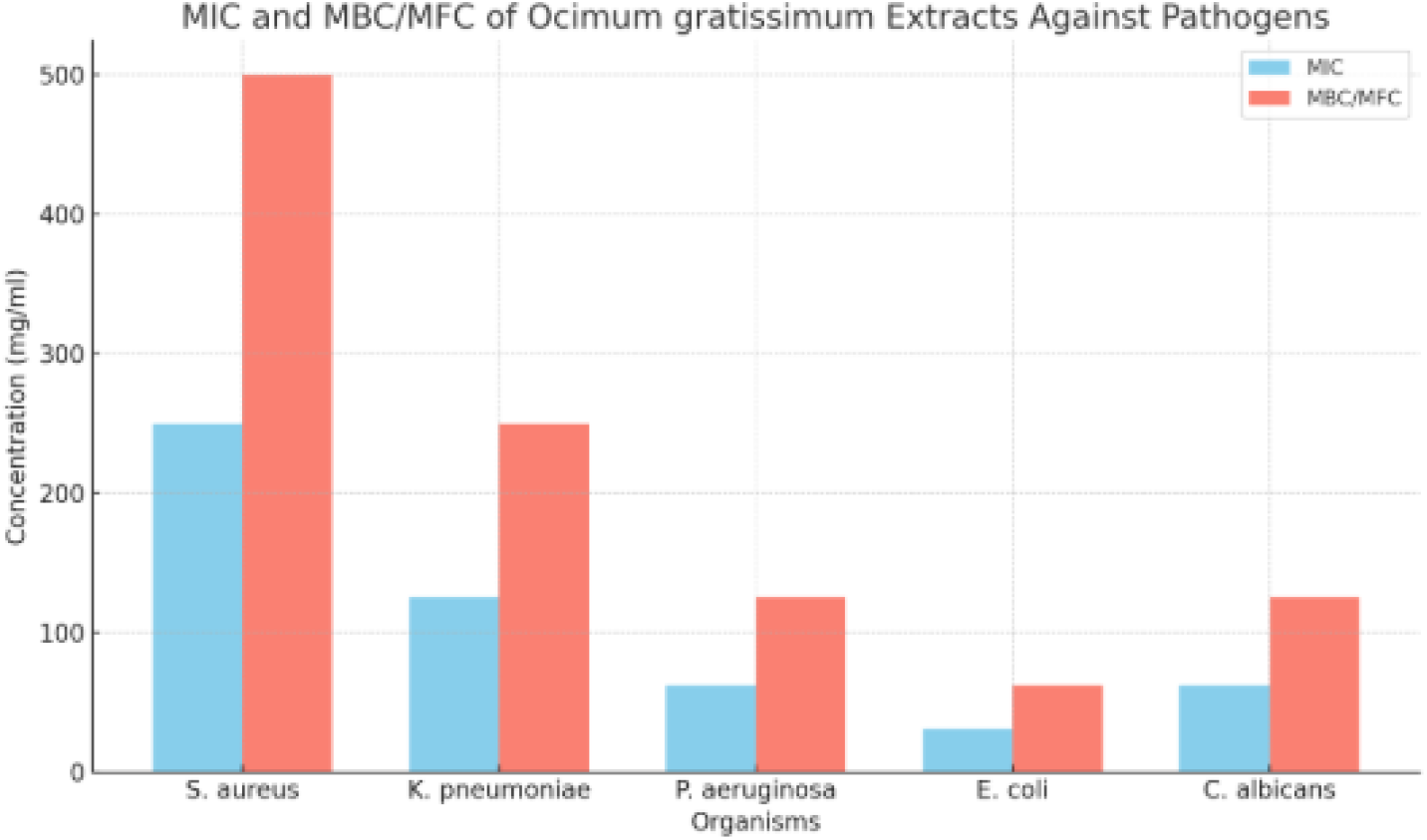
The bar chart above visually compares the Minimum Inhibitory Concentration (MIC) and Minimum Bactericidal/Fungicidal Concentration (MBC/MFC) values of *Ocimum gratissimum* extracts against various pathogens.

E. coli is the most sensitive, requiring the lowest concentrations for both inhibition and killing (MIC = 31.2 mg/ml; MBC = 62.5 mg/ml). Staphylococcus aureus is the most resistant, requiring much higher concentrations (MIC = 250 mg/ml; MBC = 500 mg/ml). Candida albicans, a fungal pathogen, also showed good susceptibility (MIC = 62.5 mg/ml; MFC = 125 mg/ml). The pattern shows that the extract possesses both bacteriostatic and bactericidal/fungicidal properties, with consistent activity across multiple pathogens.

## 6.0 DISCUSION

This study investigated the antimicrobial activity of *Ocimum gratissimum* (African basil) leaf extracts against a range of clinically significant pathogens, including *Staphylococcus aureus, Escherichia coli, Pseudomonas aeruginosa, Klebsiella pneumoniae*, and *Candida albicans*. Extracts were prepared using methanol, ethanol, and water, and their efficacy was assessed through zones of inhibition, minimum inhibitory concentration (MIC), and minimum bactericidal/fungicidal concentration (MBC/MFC). The results confirm that *O. gratissimum* possesses broad-spectrum antimicrobial properties, validating its long-standing use in traditional medicine for treating infectious diseases.

Methanolic extracts consistently showed the strongest antimicrobial effects, followed by ethanolic and then aqueous extracts. The superior efficacy of methanol is attributed to its ability to extract a broader range of phytochemicals such as eugenol, flavonoids, tannins, and thymol compounds known for their potent antimicrobial action. In contrast, the aqueous extract showed minimal activity, particularly at lower concentrations, emphasizing the role of solvent type in determining extract potency.

The antimicrobial activity was dose-dependent across all solvents. Higher concentrations produced larger inhibition zones, with the methanolic extract at 500 mg/ml yielding the highest inhibition (23.0 mm). This confirms that the antimicrobial activity of *O. gratissimum* is concentration-dependent, making it suitable for therapeutic applications when properly dosed.

Among the test organisms, *E. coli* and *C. albicans* were the most susceptible, exhibiting low MIC values of 31.2 mg/ml and 62.5 mg/ml respectively. *S. aureus*, on the other hand, showed the highest resistance, requiring a MIC of 250 mg/ml and MBC of 500 mg/ml. These differences in susceptibility could relate to structural variations in microbial cell walls and resistance mechanisms.

Findings suggest that the extract not only halts microbial proliferation but also has lethal effects on pathogens at higher concentrations. Not only did it inhibit growth, but it also killed organisms at specific concentrations *E. coli* at 62.5 mg/ml and *C. albicans* at 125 mg/ml. This dual mechanism enhances its therapeutic potential in infection management.

Furthermore, the study revealed synergistic effects when *O. gratissimum* was combined with *Vernonia amygdalina* and *Bryophyllum pinnatum*, significantly improving antimicrobial efficacy. These findings suggest that combining O. gratissimum with other botanicals may enhance its therapeutic effectiveness and reduce the need for higher doses. *O. gratissimum* is a promising natural antimicrobial agent. Its broad-spectrum activity, dose responsiveness, and synergistic potential make it a valuable candidate for developing plant-based alternatives to synthetic antimicrobials, especially in the fight against resistant infections.

## 7.0 Conclusion

This study provides clear evidence that *Ocimum gratissimum* leaf extracts possess notable antibacterial and antifungal properties, particularly when extracted with methanol. Among the tested pathogens, *Escherichia coli* and *Candida albicans* were the most susceptible, while *Staphylococcus aureus* showed the highest resistance. The antimicrobial effects were clearly concentration-dependent, with methanolic extracts consistently producing the largest zones of inhibition and the lowest MIC and MBC/MFC values. These findings support the traditional use of *O. gratissimum* for treating infections and underscore its potential as a natural alternative to synthetic antimicrobials, especially in areas facing increasing drug resistance and limited access to conventional therapies. The dual action both inhibitory and microbicidal adds to its therapeutic appeal. Future work should focus on refining extraction techniques, ensuring safety through toxicological studies, and exploring synergistic effects with other medicinal plants to improve efficacy. With further development, *O. gratissimum* could serve as a valuable component in the formulation of plant-based antimicrobial products.

### Recommendation

In light of the findings from this study, it is recommended that *Ocimum gratissimum* be further explored for the development of standardized, plant-based antimicrobial formulations, especially for treating infections caused by *Escherichia coli* and *Candida albicans*. Future research should focus on optimizing extraction using safer solvents like ethanol, conducting toxicological evaluations to ensure safety for human use, and exploring synergistic combinations with other medicinal plants to enhance efficacy. Given its demonstrated potential to inhibit and kill both bacterial and fungal pathogens, particularly in the face of rising antimicrobial resistance, *O. gratissimum* should be considered a viable alternative in natural therapy development and integrated into public health strategies, especially in resource-limited settings where access to conventional antibiotics is restricted.

## Acknowledgment

I acknowledge the support of my mentors and colleagues who reviewed this manuscript. No funding was received for this study. I am grateful to Abia State university for her support and also to Federal medical centre her assistance.

## Declaration

### Ethics approval

This study was approved by the Health Research Ethics Committee of the Federal Medical Centre, Umuahia, Abia State, Nigeria, with the approval letter reference number FMC/QEH/G.596/Vol.10/447. Written informed consent was obtained from all participants involved in the study. For participants who were minors, assent was obtained in addition to consent from their parents or legal guardians, in accordance with national ethical guidelines. The confidentiality and anonymity of all participants were strictly maintained. No identifying information, such as names of institutions or individuals, was included in the data collection or analysis. If required during the review process, a scanned copy of the ethical clearance letter will be submitted upon request.

## Authors’ contributions

Ebiloma Samuel is the sole contributor to this manuscript.

## Funding

No funding was received for this work.

Availability of data and material: Not applicable

## References

Abbara, S., Guillemot, D., Smith, D., El Oualydy, S., Kos, M., Poret, C., … Watier, L. (2024). Antimicrobial Resistance as Risk Factor for Recurrent Bacteremia after Staphylococcus aureus, Escherichia coli, or Klebsiella spp. Community-Onset Bacteremia. Emerging Infectious Diseases, 30(5), 974–983. 10.3201/eid3005.231555

Al-Rimawi, F., Sbeih, M., Amayreh, M., et al. (2024). Evaluation of the antibacterial and antifungal properties of oleuropein, Olea europaea leaf extract, and Thymus vulgaris oil. BMC Complementary Medicine and Therapies, 24, Article 297. 10.1186/s12906-024-04596-x

Chatepa, L. E. C., Mwamatope, B., Chikowe, I., Masamba, K. G., & et al. (2024). Effects of solvent extraction on the phytoconstituents and in vitro antioxidant activity properties of leaf extracts of the two selected medicinal plants from Malawi. BMC Complementary Medicine and Therapies, 24, 317. 10.1186/s12906-024-04619-7

Dantas, T. dos S., Machado, J. C. B., Ferreira, M. R. A., & Soares, L. A. L. (2025). Bioactive plant compounds as alternatives against antifungal resistance in the Candida strains. Pharmaceutics, 17(6), 687. 10.3390/pharmaceutics17060687

Dembińska, K., Shinde, A. H., Pejchalová, M., Richert, A., & Swiontek Brzezinska, M. (2025). The Application of Natural Phenolic Substances as Antimicrobial Agents in Agriculture and Food Industry. Foods, 14(11), 1893. 10.3390/foods14111893

Donkor, M.N., Donkor, AM. & Mosobil, R. (2023) Combination therapy: synergism among three plant extracts against selected pathogens. BMC Res Notes 16, 83 (2023). 10.1186/s13104-023-06354-7

Eze, A. E., Ubah, C. O., & Nwankwo, N. I. (2023). Evaluation of antimicrobial activities of some medicinal plants used in Southeastern Nigeria. African Journal of Microbial Research, 17(4), 128–137. 10.5897/AJMR2023.9921

Ezeadila, J. O., Uba, C. C., Udemezue, O. I., Ilo, P. C., Orji, C. C., & Anene, C. C. (2024). In-vitro antifungal activity of ethanol plant extracts of Moringa oleifera, Vernonia amygdalina and Ocimum gratissimum against some clinical Candida species. Microbiology Research Journal International, 34(10), 49–58. 10.9734/mrji/2024/v34i101490

Meron, G. Yonatan N. & Naimuddin, M. (2023). Phytochemical composition, antioxidant, antimicrobial, antibiofilm, and antiquorum sensing potential of methanol extract and essential oil from Acanthus polystachyus Delile (Acanthaceae). ACS Omega, 8(45), 43024–43036. 10.1021/acsomega.3c06246

Ogbolu, D. O., Oni, A. A., Daini, O. A., & Oloko, A. P. (2021). In vitro antimicrobial properties of Ocimum gratissimum extracts on bacterial isolates from infected wounds. Journal of Medicinal Plants Research, 9(6), 160–167.

Okeke, M. I., Iroegbu, C. U., & Eze, E. N. (2023). Phytochemical and antimicrobial screening of leaf extracts of some Nigerian medicinal plants. African Journal of Biotechnology, 22(2), 119–125.

Okoye, E. C., Anyanwu, M. U., & Ezeonu, C. S. (2021). Phytochemical analysis and antimicrobial activities of Ocimum gratissimum leaf extract. International Journal of Herbal Medicine, 9(3), 112–118. 10.22271/flora.2021.v9.i3b.1247

